# A cautionary tale on proper use of branch-site models to detect convergent positive selection

**DOI:** 10.1101/2021.10.26.465984

**Authors:** Amanda Kowalczyk, Maria Chikina, Nathan L Clark

**Affiliations:** University of Pittsburgh, Department of Computational and Systems Biology; University of Utah, Department of Human Genetics

## Abstract

Comparative genomics has become a powerful tool to elucidate genotype-phenotype relationships, particularly through the study of convergently acquired phenotypes. By identifying genes under positive selection specifically on branches with the convergent phenotype we can potentially link genes to that phenotype. Such gene scans are often done using branch-site codon models. However, we have observed a recent troubling trend of misinterpretation of branch-site models in which phylogeny-wide positive selection is not distinguished from positive selection specific to convergent branches. Here, we use simulations and real data to demonstrate how failing to discern between these two cases leads to false inferences of positive selection associated with a convergent trait. We then present a “ drop-out” test solution to distinguish the two cases and thereby truly capture positive selection events associated with convergent phenotypes, thus allowing for further insights into both genetic and phenotypic evolution.

## Introduction

Researchers have long used comparative biology to understand adaptations to biological challenges. One powerful comparative approach is to study independent evolutionary lineages that convergently experienced an evolutionary pressure. Traits that evolved in species sharing that pressure, but not in other species, could be inferred to be involved in an adaptive response. This strategy is now frequently applied using genome sequences from multiple species to identify genes or regulatory sequences potentially associated with their convergent response.

Multiple methods have been developed to find such genes by scanning for shifts in a gene’s overall evolutionary rate occurring preferentially in species sharing the evolutionary pressure, including RERconverge [1], Forward Genomics [2], PhyloAcc [3], and HyPhy RELAX [4]. Rather than looking for shared changes at a particular nucleotide or amino acid site, these methods identify genes that are evolving at different rates in the convergently evolving species compared to all other species. They have been applied to discover genes involved in phenotypes such as vision loss [5,6], transition from a terrestrial to a marine environment [3,7], reduction of hair density [8], and increased lifespan [9] in mammals, complex sociality in bees [10], endosymbiosis in proteobacteria [4], and loss of flight in birds [3]. Importantly, these methods have statistical designs that assure statistically significant genes experienced a rate change preferentially in the convergently evolving species. However, we have seen the rise of another approach to study convergent evolution that does not assure the same, and which could lead to false positive claims.

The concerning approach intends to identify genes that experienced positive selection specifically on branches leading to the convergently evolving species, i.e. the foreground branches. Positive selection is inferred on foreground branches when a class of codons is found to have a *d*_N_/*d*_S_ ratio significantly greater than one, which is a hallmark of positive selection [11]. Repeating this test serves as a screen for genes that responded adaptively to convergent selective pressure. While the conceptual framework behind testing for foreground-specific positive selection is logical, the interpretation of the models in many studies is fundamentally flawed.

The problem arises specifically when the tests are performed using the branch-site models in the PAML package [12] that test for *d*_N_/*d*_S_ > 1 on branches of interest. PAML is widely used in the molecular evolution community, and its branch-site models can be used to detect evidence of positive selection that acted on a particular branch or set of branches [13]. Although PAML was designed to detect positive selection on sites or branches, it may also be used to test for foreground-specific or convergent positive selection. However, it implements a limited number of phylogenetic models that some researchers attempt to coerce into answering the question of whether branch-specific positive selection exists. The typical approach is to compare the likelihood of the selection model that allows for a class of sites with *d*_N_/*d*_S_ > 1 on foreground branches to the null model that does not allow sites with *d*_N_/*d*_S_ > 1 on any branches.

The problem with using PAML to detect positive selection in response to a convergent pressure is that the results of the branch-site models are misinterpreted; while they do test for positive selection on foreground branches, they do **not** test for the **absence** of positive selection on **background** branches. A significant result from a branch-site test could indicate either that the foreground species have *d*_N_/*d*_S_ > 1 and the background species have *d*_N_/*d*_S_ ≤ 1 (**Fig. 1**) or that both foreground and background branches have *d*_N_/*d*_S_ > 1 (**Fig 1**). This is because, in the case of a gene truly experiencing positive selection on all branches, the branch-site model fits better when allowing positive selection on foreground branches, even though it would fit even better if positive selection were allowed on all branches. Therefore, **significant results from the PAML/codeml branch-site models alone do not reliably identify genes under positive selection associated with a convergent trait**. This approach runs the risk of inferring that genes evolving under positive selection on most or all branches, such as genes interacting with pathogens, are instead specifically responding to the convergent selective pressure. Unfortunately, branch-site models are frequently misinterpreted in this manner [14–19], leading to the erroneous conclusion that vast swaths of genomes are under positive selection in response to the focal convergent trait, when in fact many of those genes likely experienced tree-wide positive selection.

**Figure 1.**
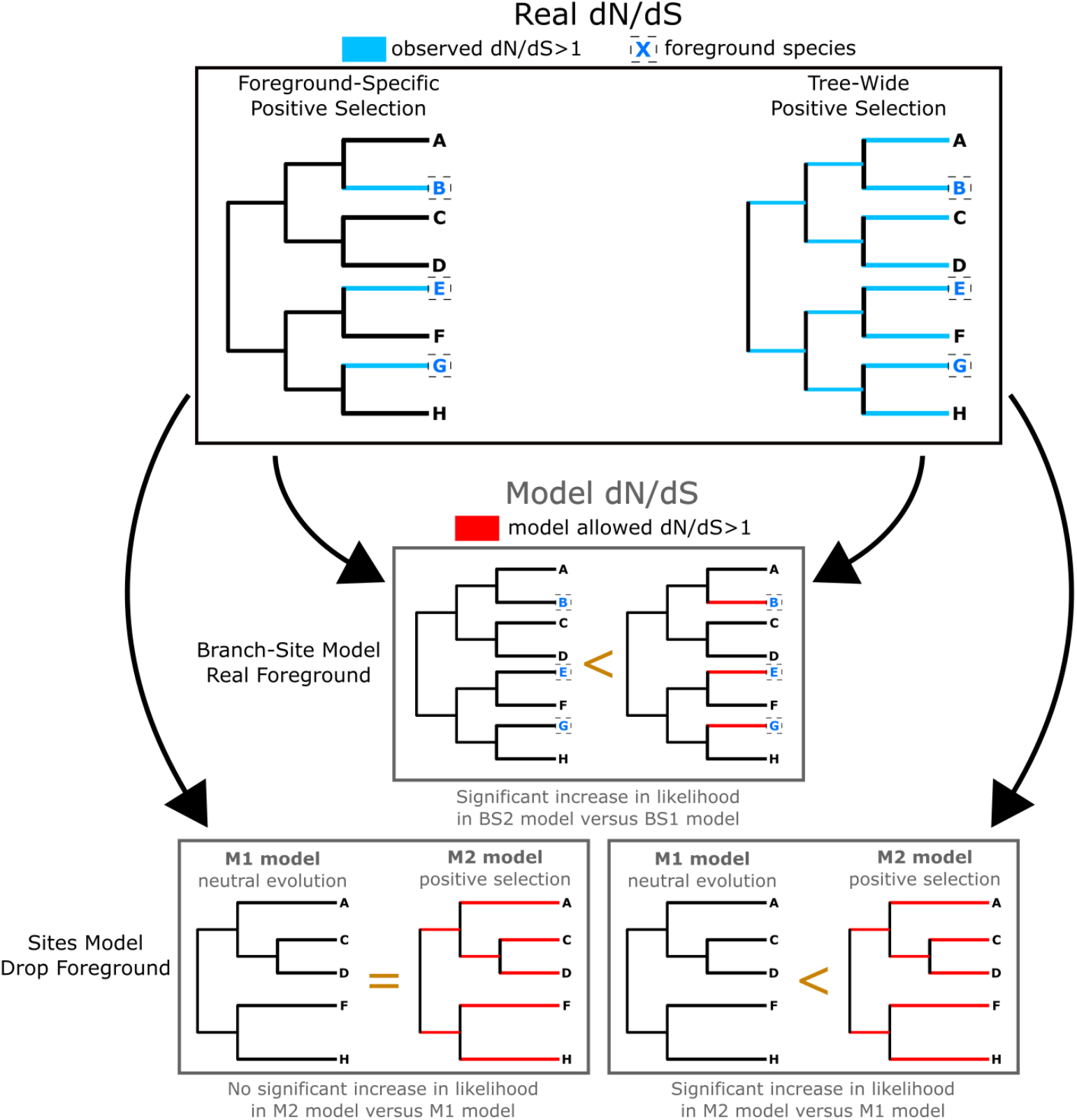
Branch-site models alone cannot detect positive selection unique to foreground branches. Consider two cases shown in the top box: the case of positive selection unique to foreground branches (Foreground-Specific Positive Selection) and the case of positive selection on all branches (Tree-Wide Positive Selection). In both cases, branch-site models for positive selection on the foreground branches will show significant signals of positive selection because there is positive selection along those branches (Branch-Site Model Real Foreground). However, if foreground branches are removed and sites models to test for tree-wide positive selection are performed, the two cases will behave differently (Sites Model Drop Foreground). In the case of foreground-specific positive selection, sites models do not show signals of tree-wide positive selection. However, in the case of tree-wide positive selection, sites models will show signals of tree-wide positive selection through the drop foreground analysis.

To illustrate, we demonstrate how misinterpretation of branch-site models leads to false inferences of positive selection in response to a convergent pressure. We then propose a simple “ drop-out” test that filters out genes with evidence for positive selection more broadly than in the foreground species. The drop-out test is a sites model test using only non-convergent species to test for tree-wide positive selection (**Fig 1**). Genes that demonstrate evidence for positive selection in species of interest from branch-site models **and** show no evidence for tree-wide positive selection are better candidates for convergent positive selection unique to foreground species. We provide code and documentation to run our recommended tests (https://github.com/nclark-lab/bsmodels-dropout).

### New Approaches

While it is possible to detect foreground specific positive selection using custom models from flexible frameworks such as HyPhy, within the context of PAML we recommend performing an additional drop-out test to confirm that positive selection is foreground specific. This can be easily accomplished within the existing PAML for branch-site analysis and we provide detailed instructions here https://github.com/nclark-lab/bsmodels-dropout. In brief, after performing branch-site tests to detect positive selection in foreground species, those foreground species should are removed and *codeml* sites models are reanalyzed. Only when the drop-out test fails to detect positive selection claims of convergent positive selection can be made.

## Results and Discussion

We present branch-site models for positive selection and drop-out site models to test for tree-wide positive selection performed on simulated data and real data. These results demonstrate that the drop-out tests are essential to distinguish convergent positive selection from tree-wide positive selection when using PAML/codeml branch-site models.

Simulated data depicted in **Fig. 2** were generated with either a class of positively selected sites specifically along foreground branches (branch-site model simulations) or a class of positively selected sites tree-wide (sites model simulations). Regardless of simulation type, all replicates showed strong signals of positive selection along foreground branches (Fig. 2: BS2-BS1 Real Foreground). Thus, using only branch-site models, it is impossible to distinguish foreground-specific from tree-wide positive selection. Only upon removing foreground branches and running sites models did it become clear that genes simulated under foreground-specific positive selection showed little signal for tree-wide positive selection while genes simulated under tree-wide positive selection showed strong signal for such selection (**Fig. 2**). We further demonstrate in **Fig. S1-S6** that the drop-out test shows the same results across a range of simulation parameter values, such as the strength of selection.

**Figure 2.**
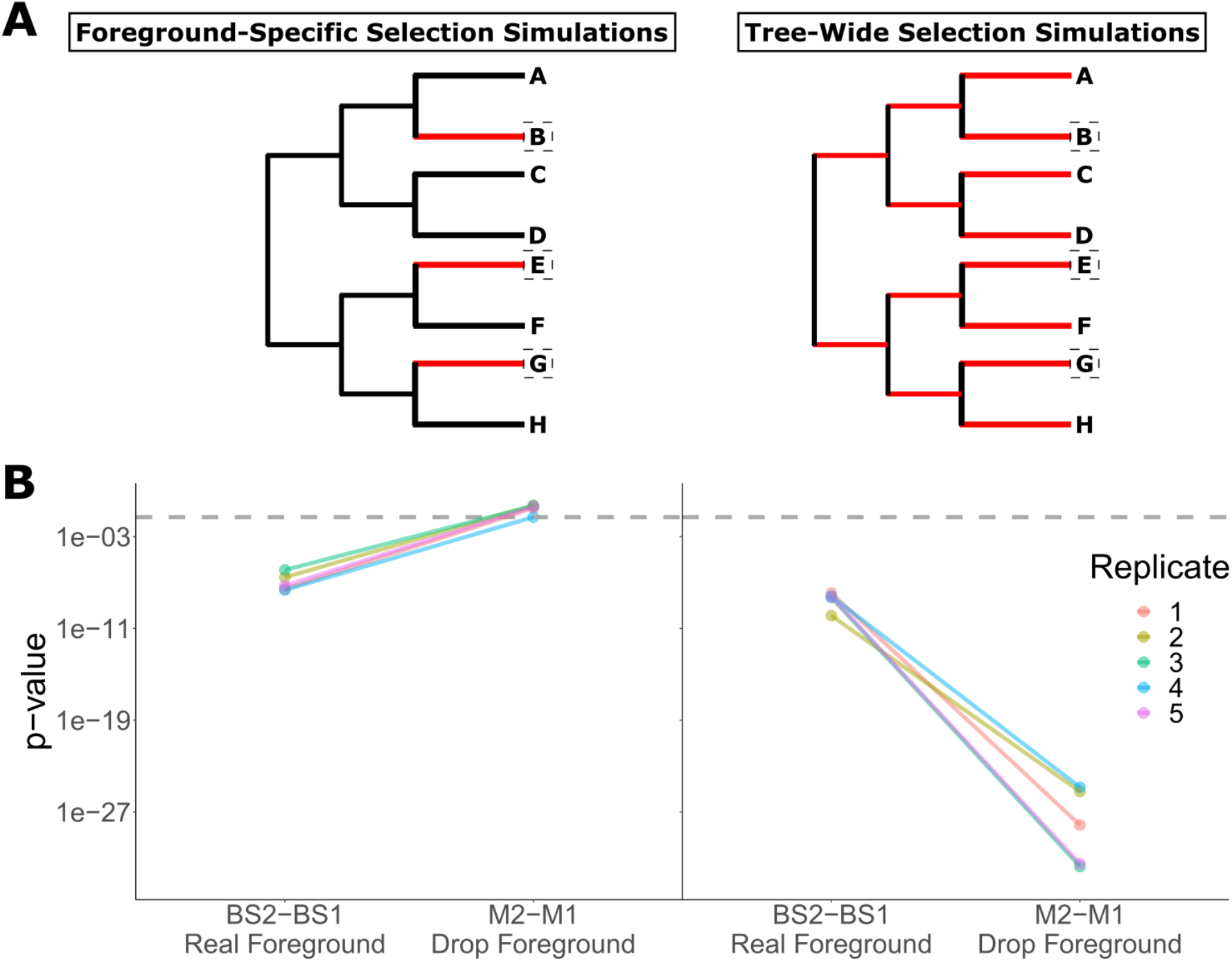
Simulated data demonstrate that branch-site models for positive selection can not distinguish positive selection unique to foreground branches from tree-wide positive selection. A) Topology of trees used to simulate sequences. In red are branches with sites under positive selection (*dN*/*dS* > 1). Boxes indicate foreground branches, which were used for branch-site models for positive selection and removed to test for tree-wide positive selection using sites models. B) Sites models distinguish foreground-specific positive selection from tree-wide positive selection. P-values from likelihood ratio tests for BS2-BS1 models (branch-site models) are indistinguishable between data simulated to have foreground-specific versus tree-wide positive selection. After foreground species are removed (Drop Foreground), p-values from M2-M1 model comparisons (sites models) are no longer significant for genes simulated with foreground-specific positive selection, while p-values remain significant for genes simulated under tree-wide positive selection, thus distinguishing the two cases. Dashed line indicates statistical significance threshold at an alpha of 0.05.

Tests on real data confirm that such concerns prevail in genuine gene sequences (**Fig. 3**). To demonstrate, we arbitrarily assigned foreground species based on a “ large” phenotype calculated from adult mass. It included 14 out of 35 mammals in the dataset, including large marine mammals, large African mammals, gorilla, and panda. Note that our test is to confirm that tree-wide positive selection can be incorrectly interpreted as foreground-specific positive selection, so the choice of foreground species is irrelevant – any set of foreground species could behave similarly. We demonstrate that four randomly selected genes with evidence of positive selection from a branch-site test also show evidence of tree-wide positive selection when foreground species are removed (**Fig. 3**). Therefore, none of the genes could be concluded to be under convergent positive selection in conjunction with the “ large” phenotype and should not be connected to its convergent evolution.

**Figure 3.**
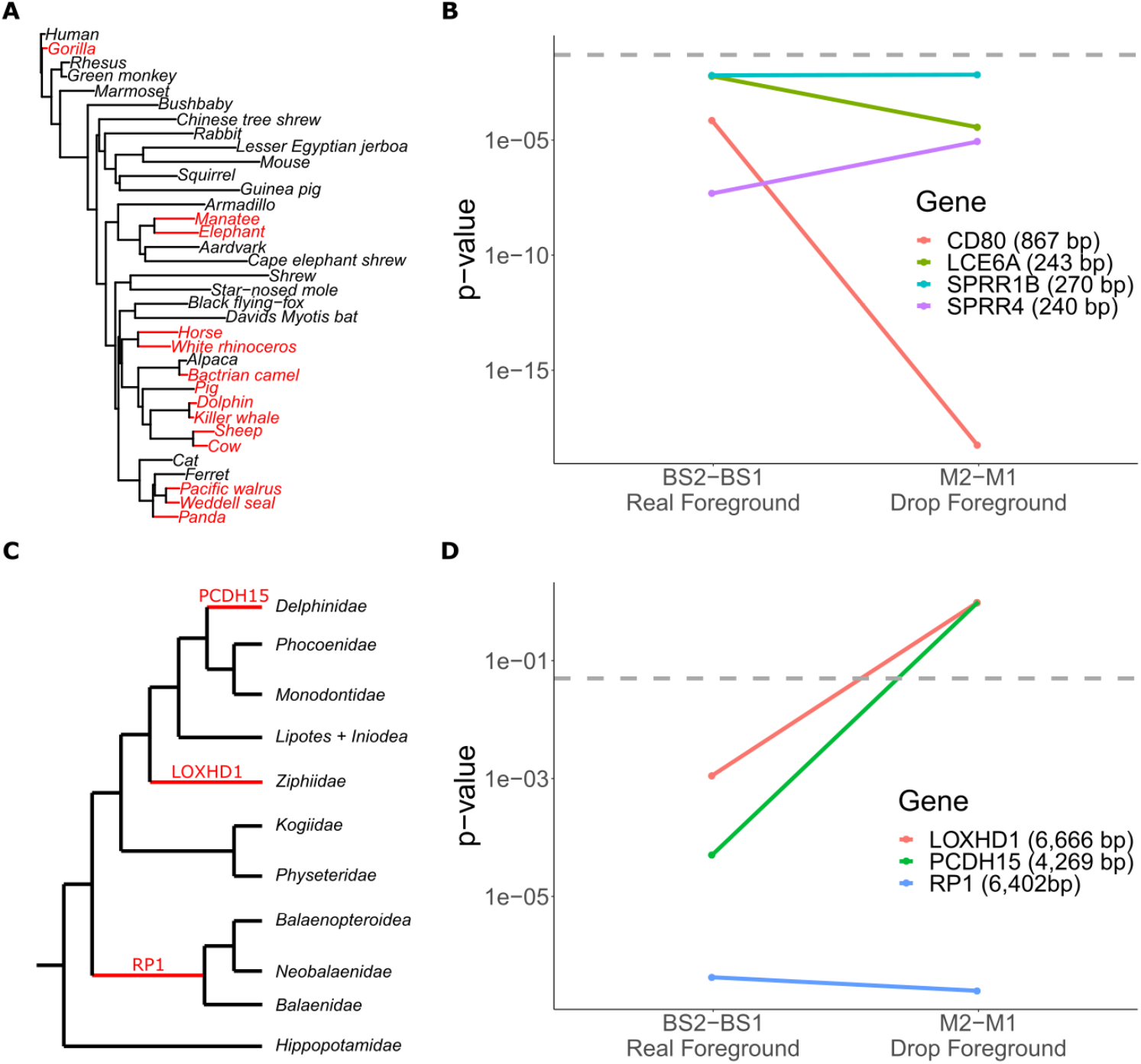
Real data demonstrate genes under foreground-specific and tree-wide positive selection in mammals. A) Full mammal phylogeny. Foreground species are labeled in red. The arbitrary phenotype “ large body size” was used solely for demonstration purposes. B) All genes with signals of positive selection from branch-site models (BS2-BS1) also show evidence of tree-wide positive selection after dropping foreground species (M2-M1). Depicted are four genes (gene length in parentheses) that demonstrate patterns of positive selection in foreground branches (according to branch-site models) and tree-wide (according to sites models). Note that without running sites models on non-foreground species, we may have falsely concluded that the genes depicted are under convergent positive selection unique to foreground species. Dashed line indicates statistical significance threshold at an alpha of 0.05. C) Phylogeny from [14] used to identify genes under positive selection in individual cetacean clades. In red are internal clade branches specified as foreground alongside the name of the gene under positive selection along that branch. D) Genes with foreground-specific positive selection in cetacean clades sometimes show tree-wide positive selection. Tests were the same as in panel B. *LOXHD1* shows positive selection in *Ziphiidae* and *PCDH15* shows positive selection in *Delphinidae* (significant result for BS2-BS1) but no patterns of tree-wide positive selection (non-significant result for M2-M1). These genes therefore show positive selection specific to their respective foreground branches. On the other hand, *RP1* shows positive selection on *Mysticeti* (significant result for BS2-BS1) and tree-wide positive selection (significant result for M2-M1), indicating that the positive selection is not specific to *Mysticeti*.

We further demonstrate that tree-wide positive selection can be misinterpreted as foreground-specific positive selection by re-analyzing data from [14] to test for positive selection on cetacean clades (**Fig. 3**). Top gene results (*LOXHD1, PCDH15*, and *RP1*) were tested for positive selection in their respective clades (*Ziphiidae, Delphinidae*, and *Mysticeti*). In this case, two of the genes did show foreground-specific positive selection (*LOXHD1* and *PCDH15*) and one (*RP1*) showed tree-wide positive selection incorrectly interpreted as foreground-specific positive selection.

We again emphasize that to identify genes under convergent positive selection unique to foreground species it is essential to perform branch-site model tests for positive selection and then filter out genes with evidence for tree-wide positive selection using drop-out tests. It is possible that studies that performed branch-site tests without drop-out tests identified genes with foreground-specific positive selection, but without a drop-out test it is unclear which genes those are. Alternatively, the authors could use one of the methods mentioned in the introduction to test for convergent rate shifts in association with their convergent trait or an additional solution presented by Davies *et al*. [20], which uses clade models to control for tree-wide positive selection [21]. Another possible strategy to control for tree-wide selection is to swap the foreground and background branches in the branch-site test. However, our simulations indicated that the swap test was less reliable than the drop-out test, and thus we recommend the drop-out test.

One limitation of our strategy is that it does not consider potential heterogeneity in positive selection throughout the tree. For example, if only a few background species experience positive selection, the drop-out test may still indicate that positive selection is foreground-specific, but at least in that case selection acted preferentially on foreground branches if not exclusively. We encourage researchers using branch-site models to investigate not only p-values, but also branch-specific omega values to identify such cases.

A formal solution to the problem we present would properly account for background selection in the model formulation. We are conducting ongoing work to implement a new likelihood model framework to properly account for background selection.

We postulate that it is unlikely to find large numbers of genes under positive selection in association with a trait because convergent positive selection is likely rare and difficult to detect outside of cases where organisms repeatedly experience a specific biochemical pressure, as exemplified by convergent drug resistance in pathogens. In previous work on various mammalian phenotypes, we indeed detected relatively little positive selection [5,7–9], suggesting that for complex phenotypes convergent positive selection strong enough to be detected by branch-site models is rare. Previous work on simulated data also suggests that branch-site tests for positive selection are conservative and are likely underpowered in cases where species are distantly-related and near synonymous mutation saturation [22]. We interpret such rarity not as a downside, but as an exciting indicator of novelty when robust signals of convergent positive selection are identified. Branch-site models are a useful tool to make such discoveries, especially when implemented with a control test to rule out phylogeny-wide positive selection.

## Materials and Methods

### Simulations

Data were simulated using *evolver* with the MCcodonNSbranchsites and MCcodonNSsites control files [12]. Note that simulated data demonstrate extreme examples of positive selection to illustrate the problem. The tree used for simulations included eight tips and the simple topology shown in **Fig. 2A**. When simulating foreground-specific positive selection, three branches spread throughout the tree were designated as foreground to imitate convergent evolution. For both sites and branch-site simulations, kappa = 1.7, # sites = 1000 codons, and tree length = 3. All non-stop-codon frequencies were set to 0.01639344 for simplicity. In sites models, three site classes were defined with omega = 0.1 at frequency = 0.15 (purifying selection), omega = 1 at frequency 0.15 (neutral evolution), and omega = 3 at frequency = 0.7 (positive selection). For branch-site models, four site classes were defined as follows: foreground and background omega = 0.1 at frequency 0.15 (uniform purifying selection), foreground and background omega = 1 at frequency 0.15 (uniform neutral evolution), foreground omega = 3 and background omega = 0.1 at frequency 0.35 (foreground positive selection, background purifying selection), and foreground omega = 3 and background omega = 1 at frequency 0.35 (foreground positive selection, background neutral). Similar distributions of omega frequencies and other parameters were used to simulate data depicted in **Fig. S1-S6**. Note that sites and branch-site simulations had the same proportion of accelerated sites either in the foreground (for branch-site models) or tree-wide (for sites models). Five replicates were generated for each type of simulation. Example control files are available at https://github.com/nclark-lab/bsmodels-dropout.

### Real Data

The mammal phylogeny was pruned from a previously reported phylogeny [23] to remove closely-related species. Such pruning reduces branch-site and sites model runtime, helps the models converge without reducing phylogenetic information, and remedies oversampling in some clades (such as primates). Mammal mass values were from the Anage Longevity Database [24], and mammals above 100,000g were defined as “ large” foreground species. Gene alignments were from the UCSC 100-way alignment [25–27]. Selected genes were pseudorandom – all genes tested had alignment file sizes equal to 15 kilobytes (a proxy for sequence length) and at least 50 mammal species of the 62 included in the full alignment. Selection of short genes allowed for fast runtimes and therefore testing of numerous genes (133 total genes) and helped ensure model convergence. Selecting highly conserved genes (present in most species) mitigated genome quality concerns and ensured that genes had similar foreground/background species counts. The four genes shown are the only genes that demonstrated foreground positive selection from branch-site models.

### Branch-site and sites models

Branch-site models were conducted using BS1 and BS2 templates with tree topologies and foreground species as shown in **Fig. 2** and **Fig. 3**. Briefly, both models specify model = 2 for two or more *dN/dS* ratios for branches and NSsites = 2 for selection. BS1 specifies fix_omega = 1 and omega = 1 to fix foreground omega at 1 while BS2 specifies fix_omega = 0 and omega = 0.4 to allow foreground sites to have omega greater than 1.

Sites models were conducted using M1 and M2 template topologies as show in **Fig. 2** and **Fig. 3** with foreground species removed. Briefly, both models specify fix_omega = 0 and omega = 0.4 to estimate omega values. M1 specifies NSsites = 1 for negative selection and neutral evolution with no codon class allowed to have omega values greater than 1. M2 specifies NSsites = 2 to build on model M1 by allowing an additional codon class with omega values greater than or equal to 1.

All parameter values are available in control files in the GitHub repository (https://github.com/nclark-lab/bsmodels-dropout).

## Acknowledgements and Funding

We thank Sergei Kosakovsky Pond for helpful discussions. This work was supported by the National Institutes of Health (R01 HG009299 and R01 EY030546 to N.C. and M.C.).

---

**Figure S1.**
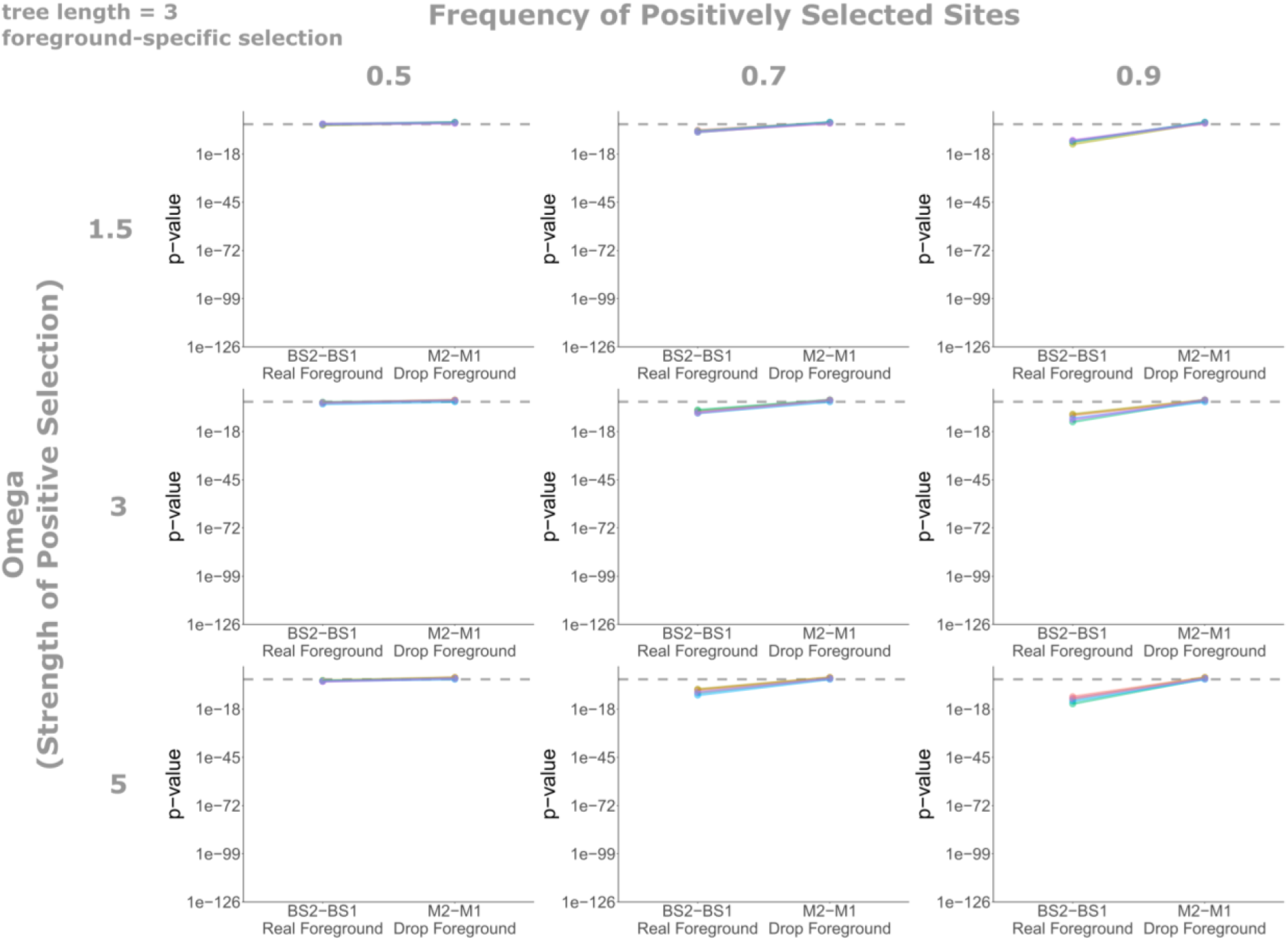
Positive selection tests on simulated data akin to those shown in Figure 2B. Data were simulated with foreground-specific positive selection, an overall tree length of 3, and multiple omegas values and frequencies of positively selected sites (see methods for details).

**Figure S2.**
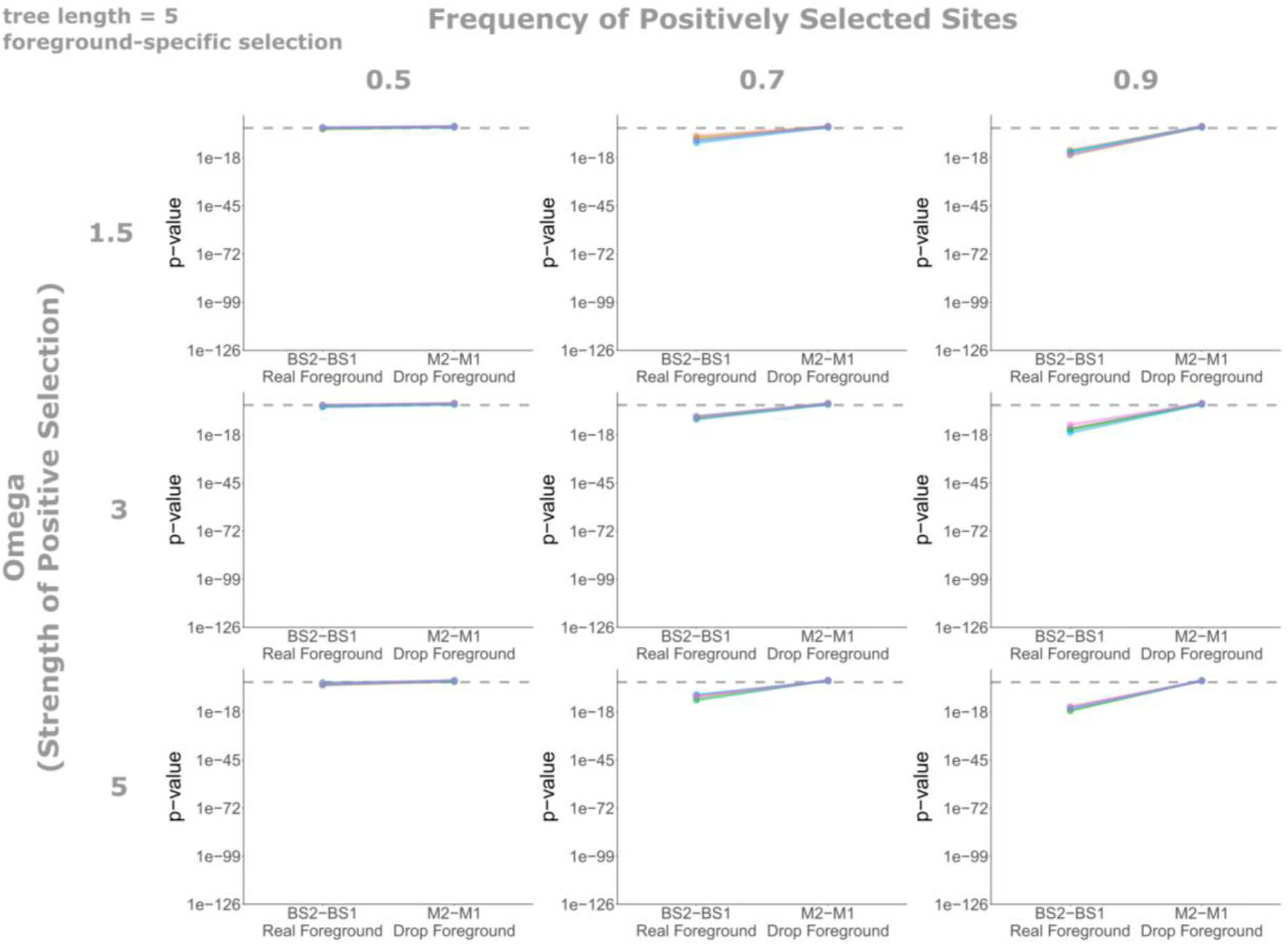
Positive selection tests on simulated data akin to those shown in Figure 2B. Data were simulated with foreground-specific positive selection, an overall tree length of 5, and multiple omegas values and frequencies of positively selected sites (see methods for details).

**Figure S3.**
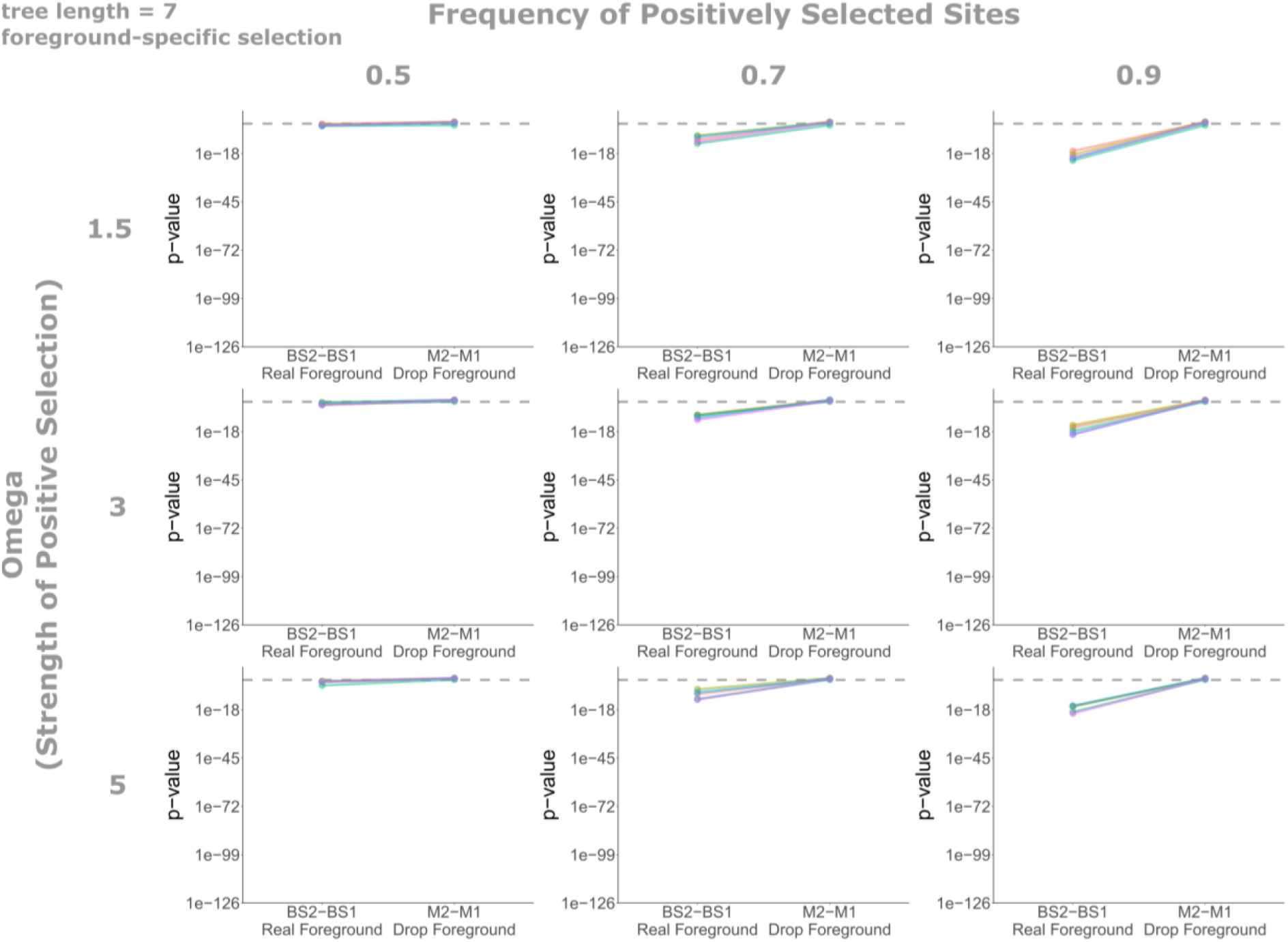
Positive selection tests on simulated data akin to those shown in Figure 2B. Data were simulated with foreground-specific positive selection, an overall tree length of 7, and multiple omegas values and frequencies of positively selected sites (see methods for details).

**Figure S4.**
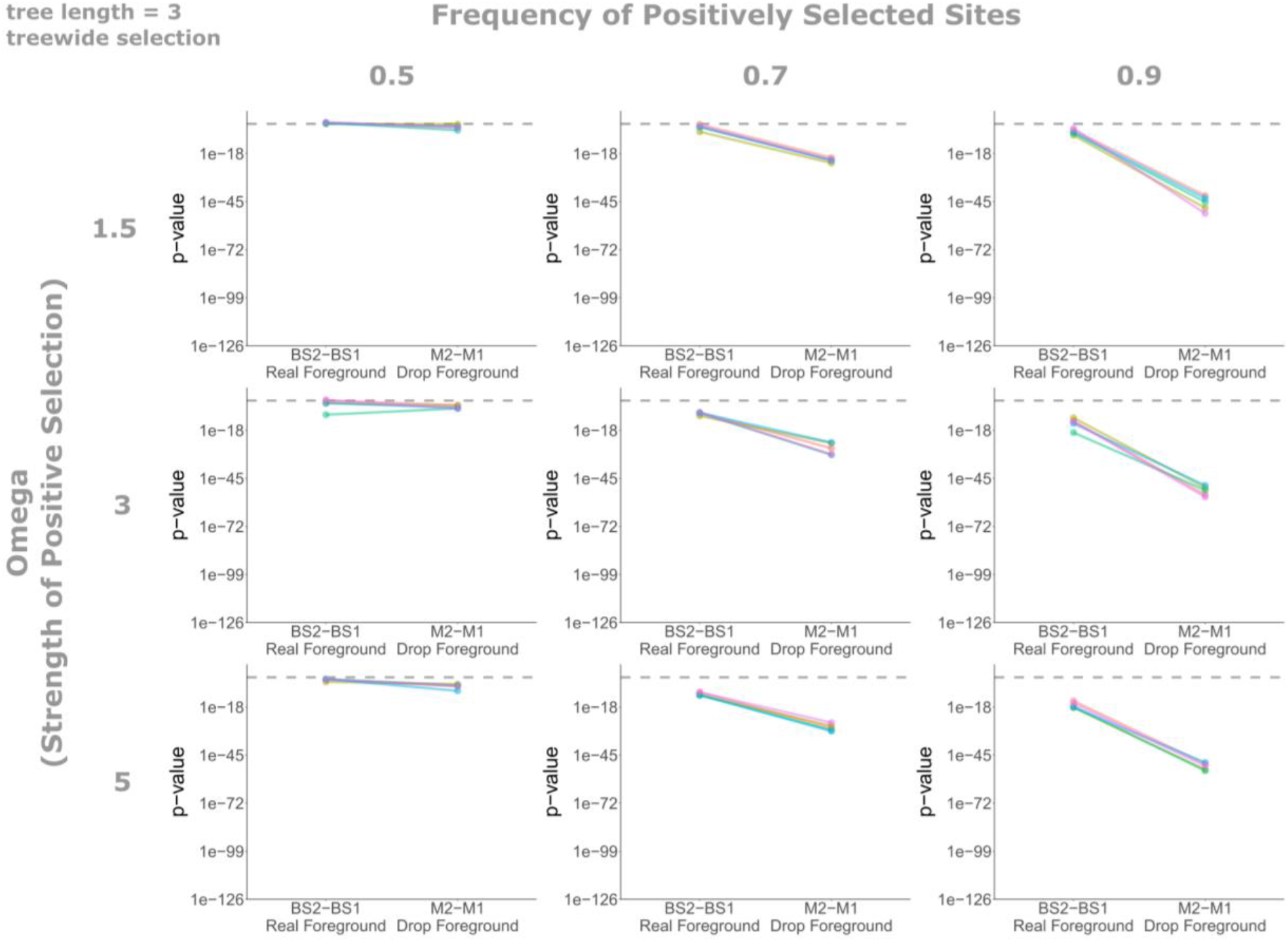
Positive selection tests on simulated data akin to those shown in Figure 2B. Data were simulated with treewide positive selection, an overall tree length of 3, and multiple omegas values and frequencies of positively selected sites (see methods for details).

**Figure S5.**
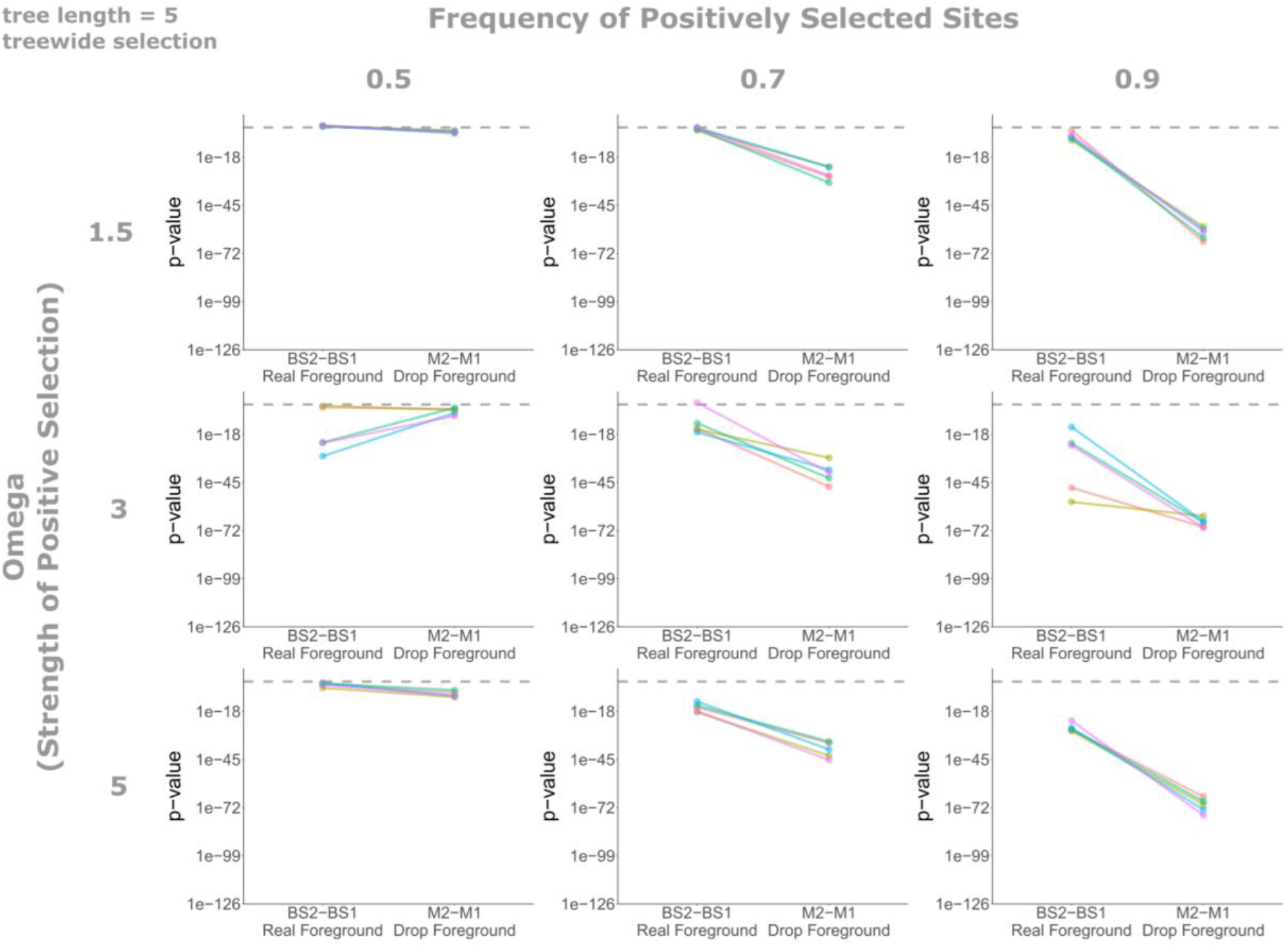
Positive selection tests on simulated data akin to those shown in Figure 2B. Data were simulated with treewide positive selection, an overall tree length of 5, and multiple omegas values and frequencies of positively selected sites (see methods for details).

**Figure S6.**
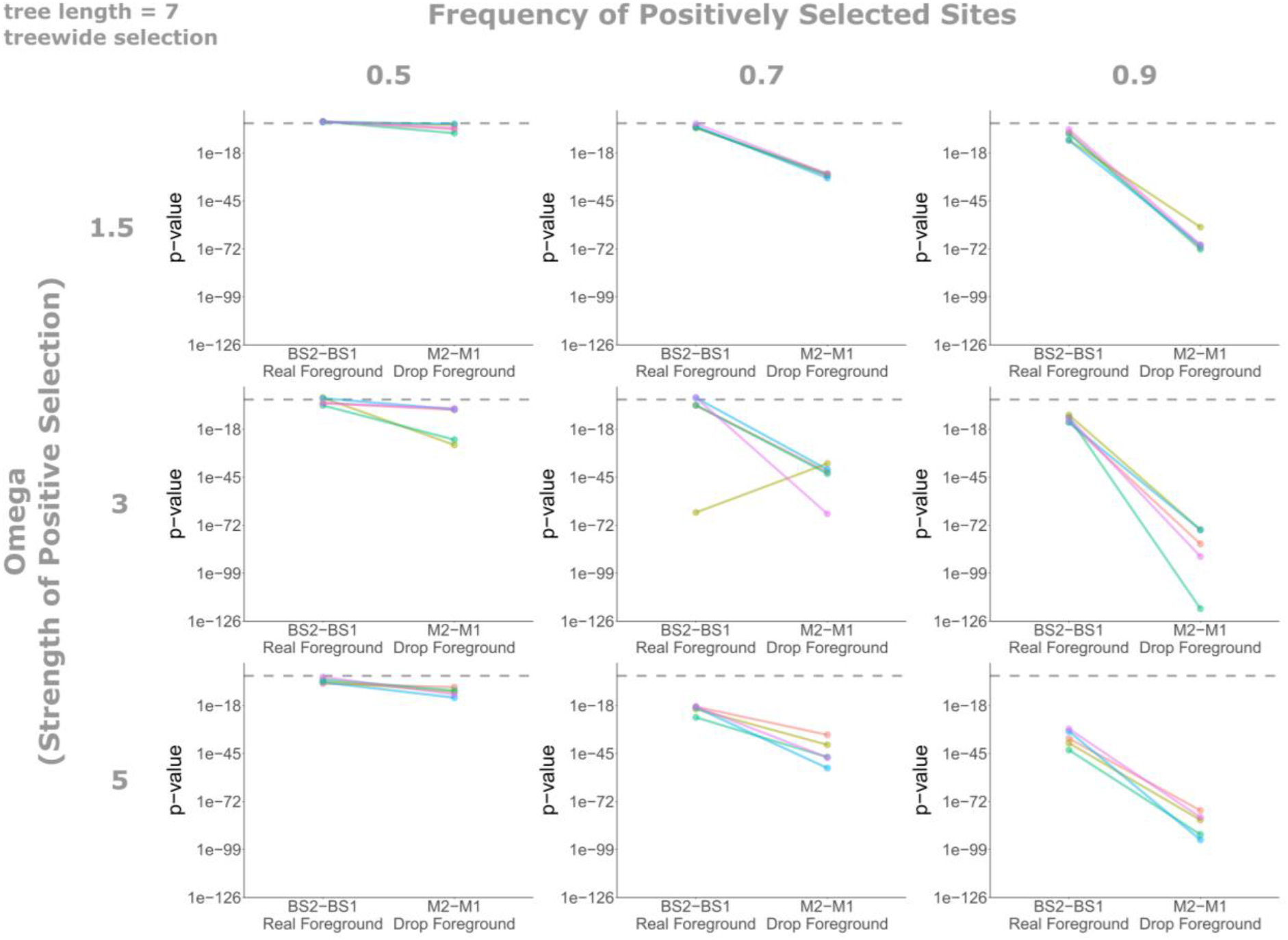
Positive selection tests on simulated data akin to those shown in Figure 2B. Data were simulated with treewide positive selection, an overall tree length of 7, and multiple omegas values and frequencies of positively selected sites (see methods for details).

